# Spatial organization of the mouse auditory cortex to sound dynamics revealed using automated image segmentation

**DOI:** 10.1101/139659

**Authors:** Ji Liu, Matthew R. Whiteway, Daniel A. Butts, Patrick O. Kanold

## Abstract

Sound stimuli are characterized by their rich spectral and temporal dynamic properties. Individual neurons in auditory cortex (ACX) encode both spectral and temporal aspects of sounds e.g. sound onset and/or offset. While the different fields of the ACX show gradients of frequency selectivity the large-scale organization of sound dynamics is unknown. We used widefield imaging of GCaMP6s in awake mouse ACX combined with a novel unsupervised image segmentation technique to investigate the spatiotemporal representation of sound onset and offset. Using this technique, we identified known auditory fields but also detected novel ACX areas. Furthermore, we found that ACX areas differed in their responses to tone onset and offset. Multiple areas were preferentially activated by tone offset, and on-response areas were more spatially localized than off-response areas. We also found tonotopy in off-responses. Together our results demonstrate a different spatial distribution of neurons across ACX for processing sound onsets versus offsets.

## Introduction

Spectral information is represented in auditory cortex (ACX) and especially in the primary auditory cortices via tonotopic maps on a scale of hundreds of micrometers, which are inherited from thalamic inputs (Guo et al., 2012; Issa et al., 2014; Kanold et al., 2014; Merzenich et al., 1975; Stiebler et al., 1997; Tsukano et al., 2015). However, it has been shown that the auditory system is not organized only according to spectral information; indeed, the dynamic nature of sound requires the auditory system to also process temporal information. ACX neurons can be sensitive to amplitude modulation, frequency modulation or sound duration (He et al., 1997; Heil et al., 1992; Schreiner and Urbas, 1986). A systematic organization with regard to frequency sweep rate has been identified in mouse ACX (Issa et al., 2017) while a map of periodicity has been proposed in cat ACX (Langner et al., 2009). At other levels of the auditory pathway such as the inferior colliculus, topographic organizations of amplitude modulation (Heil et al., 1995; Langner et al., 2002; Schreiner and Langner, 1988) and frequency modulation (Hage and Ehret, 2003) exist. Thus, the auditory system might not only be organized according to the sound frequency but also to the dynamic properties of sound.

Sound onset and offset also constitute dynamic aspects of sound, besides frequency and amplitude modulation. Indeed, neurons at multiple levels in the auditory pathway respond to sound onset and offset (He et al., 1997; Henry, 1985; Hillyard and Picton, 1978; Kopp-Scheinpflug et al., 2011) including the ACX (Baba et al., 2016; Fishman and Steinschneider, 2009; He, 2001; Qin et al., 2007; Recanzone, 2000; Scholl et al., 2010). While off-responses have been suggested to be responsible for duration coding (He, 2001), they more fundamentally reflect the auditory system’s ability to encode the sudden termination of auditory stimuli, a sharp contrast between the presence of sound and silence.

On- and off-responses are suggested to be conveyed by non-overlapping synaptic circuits (Scholl et al., 2010), raising the question whether there would be difference in the spatial representation of on- and off-response. Widefield imaging of flavoprotein signal in ACX of anesthetized mice suggested the presence of an distinct area sensitive for sound offsets, and that off-response tonotopy was absent (Baba et al., 2016). However, off-responses are most prominent in awake animals (Fishman and Steinschneider, 2009; Joachimsthaler et al., 2014; Qin et al., 2007; Recanzone, 2000). Thus, we investigated the spatiotemporal representation of tone offset in ACX in awake animals using widefield imaging of GCaMP6s.

Recent optical studies provided detailed descriptions of the organization of ACX and primarily A1 on the macro- and meso-scale level in mice (Bandyopadhyay et al., 2010; Issa et al., 2014; Issa et al., 2017; Rothschild et al., 2010; Tsukano et al., 2015; Winkowski and Kanold, 2013). Conventionally the segmentation of ACX into regions of interest (ROIs) is based on snapshots of activity following stimulus presentations, thus capturing only the on-responses. We here used the entire image sequence acquired when tones were successively played (thus capturing both on- and off-response) and developed a novel, unbiased and unsupervised method to define ROIs using a constrained latent variable model (Whiteway and Butts, 2017). This model defines ROIs based solely on the co-activation of pixels over time, which includes both periods of spontaneous and stimulus-driven activities. Conceptually, this procedure produces a lower dimensional segmentation of the image sequence, and thus aids in the understanding of both the spatial and temporal activation pattern. Using this approach, we detected previously known areas (e.g. A1) but also revealed novel auditory fields. We found that both on- and off-responses show tonotopic organization. Additionally, we found that the relative amplitude of on/off-response is not only a function on particular stimulus parameters (i.e., frequency and sound level) but also depends on particular auditory field with some field showing weak off-response while others were dominated by off-response when responding to the same tone. This suggest that different auditory fields might have different roles in temporal processing. We also found that off-response is more extensive in space than on-response, suggesting that the underlying circuits differ from those carrying on-responses and have a greater spatial extent. Our results demonstrate the existence of tonotopy in off-responses and the spatial diversity of on/off-response _5_patterns in ACX, and illustrate that on large-scales ACX is organized not only with respect to sound frequency but also with respect to the temporal aspects of the stimulus.

## Results

Neurons in ACX can respond to the onset and/or offset of sound (Baba et al., 2016; Fishman and Steinschneider, 2009; Qin et al., 2007; Recanzone, 2000; Scholl et al., 2010). To identify areal differences in on- and off-responses, we presented 2-second duration pure tones to awake adult F1 (n=13) mice from CBA/CaJ and Thy1-GCaMP6s (C57/BL6 background) crosses (Dana et al., 2014). Adult F1 CBA/CaJ x C57/BL6 mice have hearing comparable to adult CBA/CaJ mice (Frisina et al., 2011); thus our cross allows expression of GCaMP6s uniformly in A1 without the hearing loss present in adult C57/Bl6 mice. To identify auditory responsive regions, we performed widefield imaging over a cranial window of ~3mm radius over the left ACX while the mice were passively listening to tones.

Tone onset resulted in spatially restricted fluorescence increases at several locations in the imaging field (Figure 1A, see 0.4s following tone onset). Fluorescence increases were present in discrete locations corresponding to activation of putative A1, AAF and A2 respectively. Following tone offset, we observed an additional increase of fluorescence (at 2.4s, or 0.4s after tone offset), which corresponded to an off-response. On- and off-responses were also present in response to ultrasonic frequencies such as 61.3 kHz (Figure 1B). In both examples, the spatial locations of the fluorescence increase qualitatively match prior studies (Baba et al., 2016; Issa et al., 2014; Tsukano et al., 2015).

**Figure 1.**
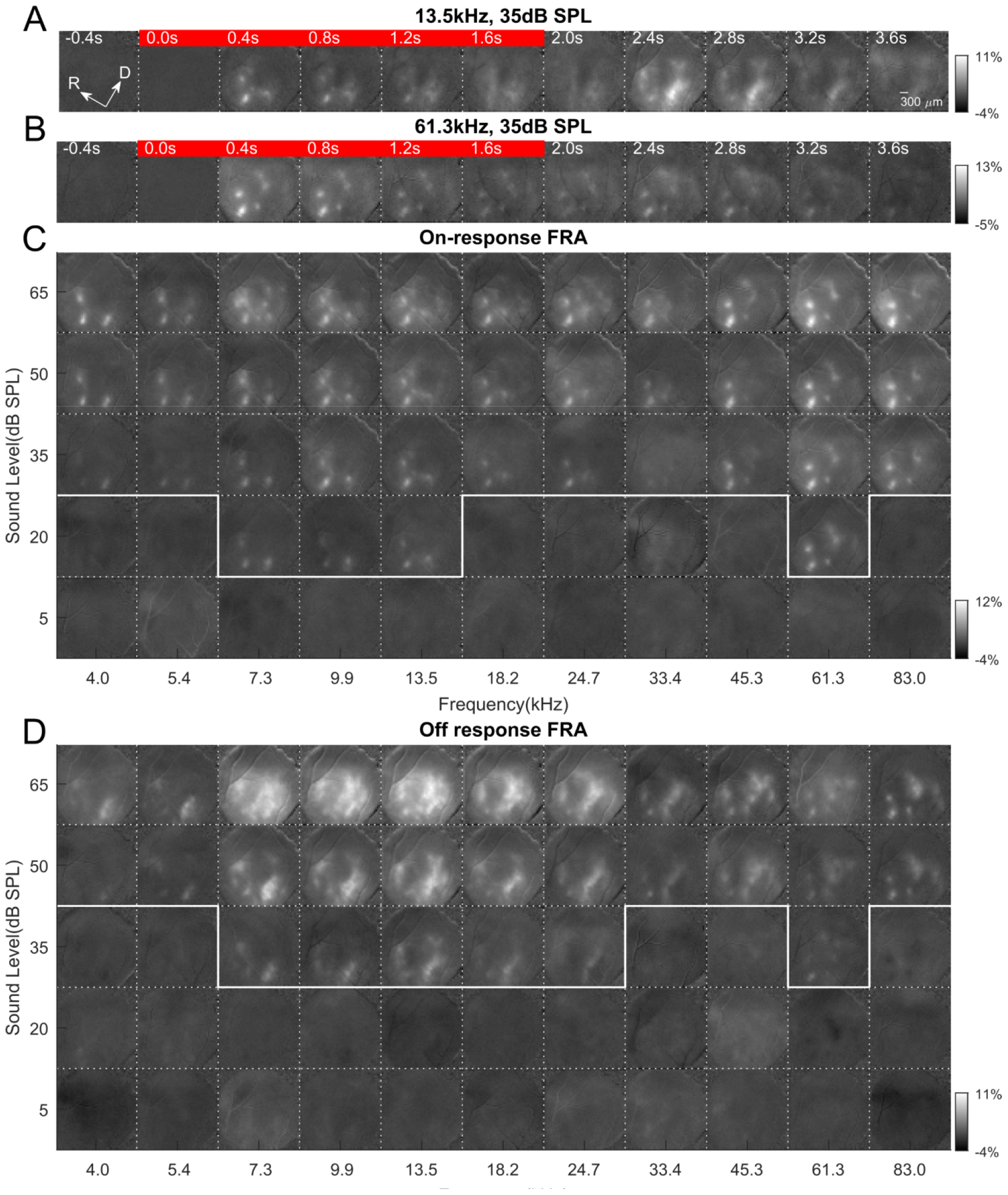
Widefield example image sequence and on/off-response FRA. (A) Sequence of widefield images showing response to 13.5kHz tone at 35dB SPL. The red bar indicates the images collected during tone presentation (0-2sec). (B) Same as in (A) but shows image sequence in response to 61.3kHz tone at 35dB SPL. (C) On-response FRA. Baseline subtracted average images within 200-500ms after tone onset are plotted as a function of frequency and sound level. White solid lines show threshold at each frequency. (D) Off-response FRA. Average images within 200-500ms after tone offset are plotted with images 0-200ms before tone offset used as baseline. Typically, off-response had a higher threshold than on-response.

To demonstrate the frequency and sound level dependence of the widefield on/off-response, we tiled snapshots of activities following tone onset or offset (Figure 1C, D). We noted that on-responses were generally evoked at lower sound levels than off-responses. In addition, both the spatial locations of on-responding areas and off-responding areas seemed to show systematic changes with respect to sound frequency. Moreover, while on-responses occurred in spatially restricted areas, off-responses appeared to occur in larger and more diffusive areas. To quantify these observations in detail, we performed image segmentation to define distinct ROIs and to obtain their temporal activation profiles.

### Unbiased and unsupervised detection of ROIs

Prior studies have used widefield imaging to calculate stimulus sensitivity for individual pixels (Issa et al., 2014; Juavinett et al., 2017). However, these sensitivity maps usually reflect cortical organization with respect to single stimulus properties, e.g. frequency and retinal location. Here, we aim to determine stimulus sensitivity for both on- and off-response at the same time. Using prior approaches would generate two separate maps, which could be difficult interpret. Alternatively, ROIs can be defined such that on- and off-response of the same spatial area can be compared. To achieve this goal, we developed an unbiased and unsupervised image segmentation technique to define ROIs. The goal of our image segmentation is to use dimension reduction techniques to break down the image sequence into linear combinations of ROIs weighted by their respective activity (Figure 2A).

**Figure 2.**
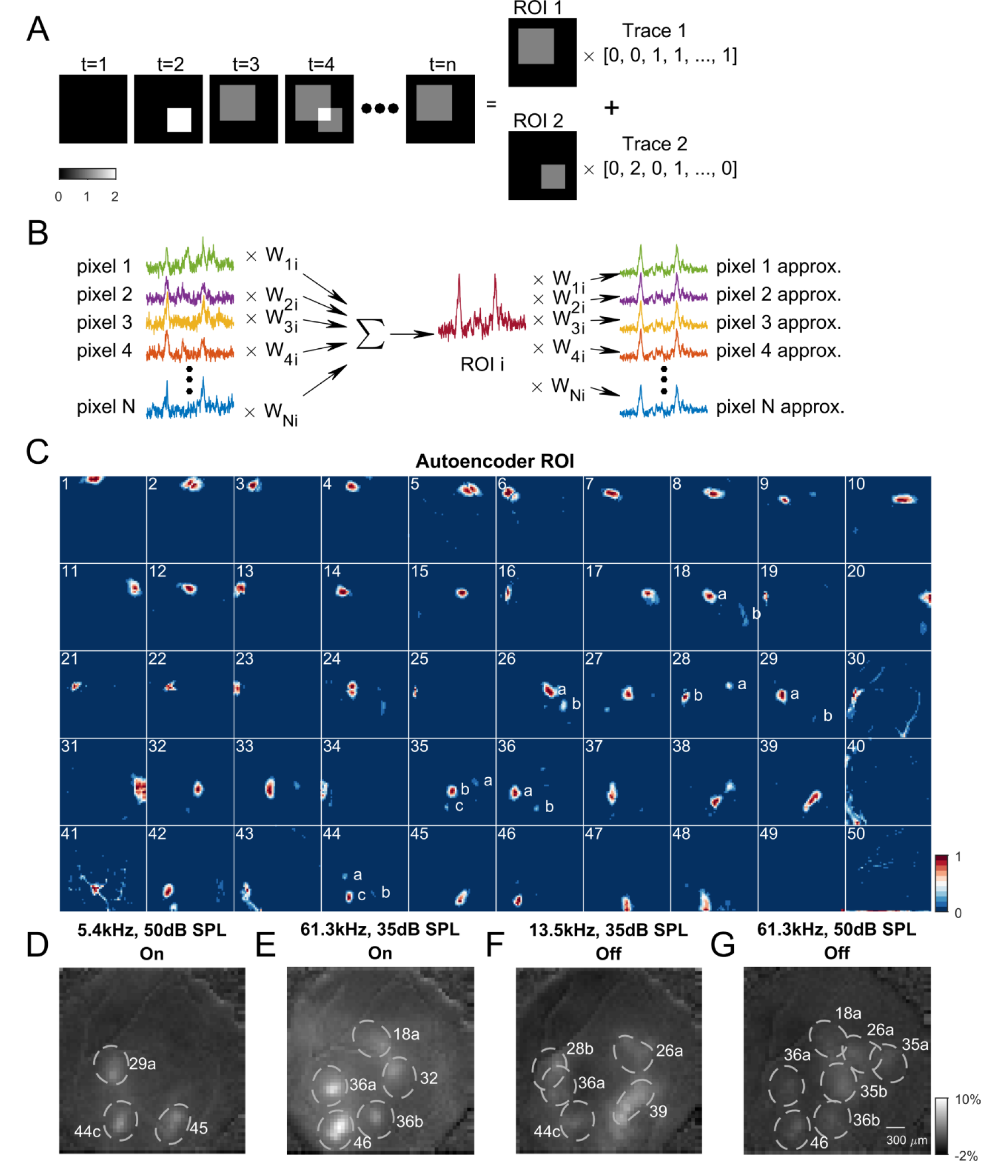
Autoencoder was used to perform widefield image segmentation. (A) Cartoon showing image segmentation. The example image sequence at any time point can be expressed as the weighted summation of ROI 1 and ROI 2 by respective activity level. Our goal of image segmentation is thus to retrieve activated areas as well as their temporal activation traces. (B) Principle of fitting autoencoder ROIs. Temporal trace of *i*th ROI is constructed by linearly weighting all pixels at each time point. Weights from *j*th pixel to *i*th ROI is *W_ij_*. Weights are adjusted such that *i*th ROI’s trace, when reciprocally weighted by *W_ij_* and subtracted from jth pixel’s trace, results in minimal error summed over all pixels. *W_ij_* forms *i*th ROI’s spatial profile.(C) Autoencoder ROIs fitted on same widefield images as in Figure 1. Note that some ROIs have more than one distinct spatial areas (labeled with letters). For example, ROI 36 captures the increase in fluorescence to high frequency tones (see Figure 1B for comparison) located at presumptive AAF and A1. Also note that ROI 39 captures the elongated shape of the increase in fluorescence. (D)-(G) On- and off-response spatial profiles overlaid with selected Autoencoder ROIs to visually validate ROI placement. D-G share the same colorbar.

To perform unsupervised image segmentation, we used an autoencoder neural network with non-negativity constraints (Whiteway and Butts, 2017). An autoencoder is a neural network where the input and output layers have the same number of nodes, with one or more hidden layers between them (Figure 2B). The goal of this constrained autoencoder was to adjust the weights between the input layer and the hidden layer and those between the hidden layer and the output layer such that the output matched the input as closely as possible, while constraining the weights between the hidden and output layers to be non-negative. Because the number of nodes in the hidden layer was much smaller than the number of nodes in the input/output layers (corresponding to the number of image pixels), using this method results in a dimensionality reduction of the original image sequence. In this sense, the autoencoder is very similar to principal component analysis (PCA). However, the segmentation achieved by the autoencoder was much different than PCA (Figure S2A), because the weights between the input/output layers and the hidden layer were constrained to be non-negative, in order to ensure that any increase or decrease in fluorescence would be reflected in the temporal activity trace of the ROI without ambiguity. Thus, the temporal activation of ROIs was defined by the activity of the hidden layer, and the weights between the hidden layer and the output layer defined the coupling to each pixel of each ROI. Both the activities of the ROIs and the spatial weights were fit using the sequence of images over the entire experiment.

Typically, an autoencoder with around 50 ROIs achieved a good fit of the image sequence (Figure S1A), and the resulting ROIs densely tiled the imaged area (limited by the cranial window), indicating that most pixels have been incorporated into ROIs (Figure S1B, D). Also, these ROIs had minimum spatial overlap with each other, as shown by the spatial correlation matrix that closely resembles an identity matrix (Figure S1C). Thus, the ROIs defined spatially unique regions. While most ROIs detected in this way contained a continuous region in space (e.g., Figure 2C, ROI 1-8), some of the ROIs had more than one distinct spatial areas (e.g., Figure 2C, ROI 36). A careful examination of such ROIs show that they captured multiple regions that were co-active to the same stimuli. To ensure that ROIs represented continuous areas we performed automated split of such ROIs (see Methods).

Overlaying selected ROIs with the snapshots of activity from Figure 1C, D shows that the placement of ROIs agreed visually with the location of activation for both on-(Figure 2D, E) and off-response (Figure 2F, G), and their shapes also reflected the contour of fluorescence increase. These results show that using an autoencoder for image segmentation can produce spatially localized ROIs that tile ACX to facilitate auditory field identification, and that the defined ROIs faithfully capture the spatial activation pattern.

### Automatically identified ROIs reliably identify core ACX fields across animals

ACX of mice contains several auditory fields, including A1, AAF and Ultrasonic Field (UF), which are characterized by the presence of tonotopic gradients in the on-response (Stiebler et al., 1997). Tonotopy also exists in secondary area A2, albeit on a compressed scale (Issa et al., 2014). Having unbiasedly defined ROIs, we next sought to assign them to different auditory fields according to the frequency selectivity in on-responses and their relative spatial locations.

First, we identified A1 and UF ROIs based on their two tonotopic axes, one from the caudal side to dorsomedial side (low to high) and the other one, sharing the same low frequency area, from caudal to ventrolateral side (Issa et al., 2014). The spatial locations of example A1 and UF ROIs (Figure 3A) as well as the temporal traces as a function of frequency and sound level (Figure 3B-F) show progression of frequency selectivity along the two tonotopic axes. The most caudal ROI showed the highest on-response amplitude to low frequency tones (Figure 3B), while as one moves dorsomedially, ROI’s frequency selectivity shifts towards mid frequency range (Figure 3C, D). UF ROIs were identified dorsally located to the mid-frequency A1 ROIs (Figure 3F-H), which showed selectivity to high frequency such as 61.3kHz. A similarly high-frequency selective ROI can be identified ventral to mid-frequency A1, and we assigned this ROI to high-frequency A1 (Figure 3E). We use ‘UF’ and ‘high A1’ to distinguish between the two spatially distinct areas that are high frequency selective, while they are both considered primary auditory cortices. So far, we showed that the ROIs given by our unbiased image segmentation technique can robustly identify the two known tonotopy axes in A1, replicating the results of prior studies (Issa et al., 2014; Polley et al., 2007; Tsukano et al., 2015). Similarly, we identified AAF ROIs and recovered its tonotopy that ran from rostral side towards ventrolateral side (Figure S3B-D). We also identified A2 (Figure S3E, F), which were located most ventrolaterally and with very broad frequency tuning.

**Figure 3.**
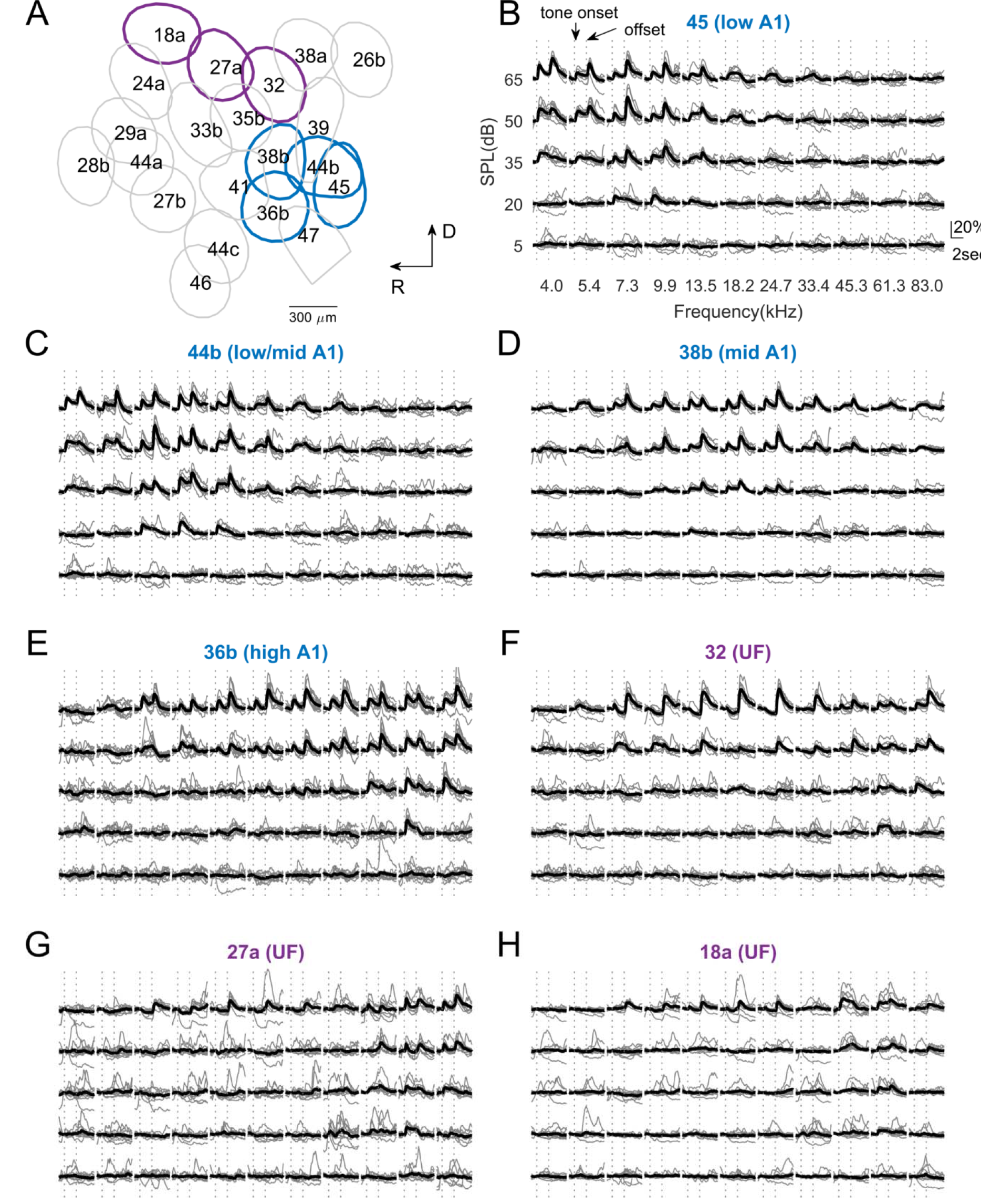
FRAs of selected ROIs from Figure 2C. (A) Overlay of selected ROIs. The number and letter correspond to ROIs shown in Figure 2B. (B-H) FRAs of selected ROIs. Axes label and y-axis scale are the same, and thus omitted from all but panel (B) for figure compactness. Paired dotted lines indicate tone onset and offset respectively. (B) low-frequency A1 ROI. (C) low/mid-frequency A1 ROI, note the smaller on-response to 4kHz tone at 35dB SPL compared to (A), and yet stronger on-response at 13.5kHz at 20dB SPL. (D) mid-frequency A1 ROI. (E) high-frequency A1 ROI. (F-H) UF ROIs.

We performed parcellation of ROIs in all animals studied, and the similar spatial layout of A1, UF, AAF and A2 can be robustly observed (Figure 4A, B). To quantify the frequency selectivity of these auditory fields, we calculated on-response amplitude and summed over sound level to obtain the onset frequency response profile (Figure 4C-G). Onset frequency response profiles for low/mid/high A1 and UF show peaks at respective frequency range (Figure 4C, D), and the same can be observed of low/mid/high AAF (Figure 4E). For A2, we separate the ROIs into two groups (low/mid and high) due to their broad tuning (Figure 4F). Here we show that known auditory fields can be identified using our novel image segmentation technique. Moreover, our results show that the spatial arrangement of known auditory fields is fairly stereotypical across animals.

**Figure 4.**
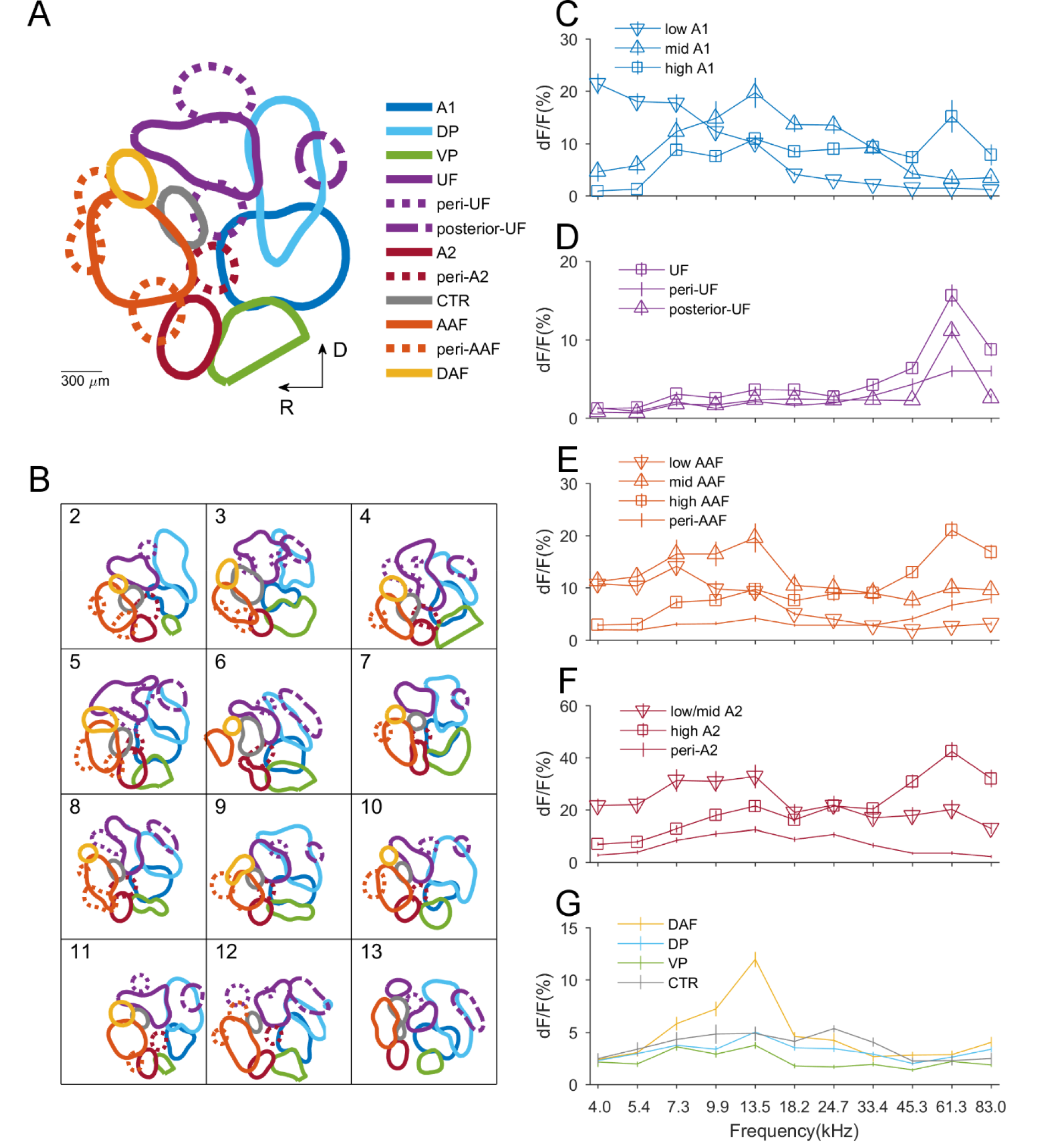
Field parcellation. (A) Same example as in Figure 1-3. ROIs are assigned to different auditory fields based on their FRAs. (B) Field parcellation results from 12 other mice in this study. (C-G) Onset frequency response profiles for different auditory fields were obtained by summing over sound level in on-response FRAs and averaging across ROIs. All errorbars show SEM.

### Automatically identified ROIs reveal novel auditory areas

The above identified areas only account for a proportion of the total ROIs, with the rest of the ROIs still capturing meaningful sound driven responses in the image sequence. This suggests that these ROIs also identify auditory areas. First, we found a subset of ROIs located dorsally to A1 which we assigned to DP (Figure S3G, H). They showed relatively weak on-responses and no prominent peak in their onset frequency response profile (Figure 4G). However, they showed prominent off-responses. We also identified a Ventral-Posterior field (VP) ventral to A1 but with similar properties as DP (Figure 4G, Figure S3I).

Caudal to DP, we could identify an area with high frequency selectivity, which we call ‘posterior-UF’ (Figure 4D, see also Figure S3J). This area is separated from UF by DP and yet its onset frequency response profile shows prominent peak at 61.3kHz. To our knowledge, it is the first time that such auditory field is identified. We also found ROIs flanking UF with similar frequency selectivity but smaller on-response (Figure 4D, Figure S3K). We assign these ROIs to ‘peri-UF’.

Dorsal to AAF, an area was found that had onset frequency response profile peaking at mid frequency, which we call Dorsal Anterior Field (DAF) (Figure 4E, see also Figure S3L). The location and frequency selectivity of DAF did not comply with the tonotopic gradient of AAF and thus was assigned a separate auditory field. We also found ROIs flanking core AAF areas which showed weaker on-responses and were assigned to ‘peri-AAF’ (Figure 4E, see also Figure S3M).

Dorsal to low/mid A2, an area was found with on-responses mostly to mid-range frequencies (Figure 4F, see also Figure S3N). We assign this area to a separate field, i.e., ‘peri-A2’, as the area’s frequency selectivity was in the opposite direction of A2 tonotopic gradient. Lastly, sandwiched between AAF and A1 we found an area that showed little on-or off-responses, and we named it Center (‘CTR’, Figure 4G, Figure S3O).

So far, our unbiased image segmentation technique has allowed a comprehensive parcellation of ACX. We have shown the identification of already known auditory field, such as A1, UF, AAF and A2. We also identified novel auditory fields, such as DAF and posterior-UF, despite their small yet consistent on-responses. Thus, we have shown that our image segmentation technique is a very sensitive and robust method to identify and delineate sensory responsive areas.

### Off-responses show tonotopic organization

We have partially established the existence of a tonotopic map for on-response using our image segmentation (Figure 4C-F). To identify if such a tonotopic organization also existed for off-responses, we selected a subset of ROIs that exhibited robust on- or off-responses at the respective threshold for different frequencies (Figure 1C, D, see white outline) and plotted the characteristic frequency determined at threshold (Figure 5). For on-responses (Figure 5A, B), we used A1 (low, mid, high), UF, AAF (low, mid, high), DAF and A2 (low/mid, high) ROIs. For off-response (Figure 5C, D) we added DP, VP and peri-AAF ROIs as they showed significant off-responses. First, we confirmed robust tonotopy for on-response in A1, AAF and A2 (Figure 5A, B), with a pattern largely consistent with prior studies (Issa et al., 2014; Tsukano et al., 2015). A subtle difference is that our results show that the two on-tonotopic gradients constitute by A1 and UF share the low to mid frequency axis. In terms of off-response, we did observe tonotopy in A1, UF, as well as in AAF and A2 in all animals studied (Figure 5C, D). The off-tonotopic gradient from A1 to UF overlapped with the on-tonotopy gradient, but it also had an additional gradient extending through DP towards UF, due to DP’s recruitment in the off-response.

**Figure 5.**
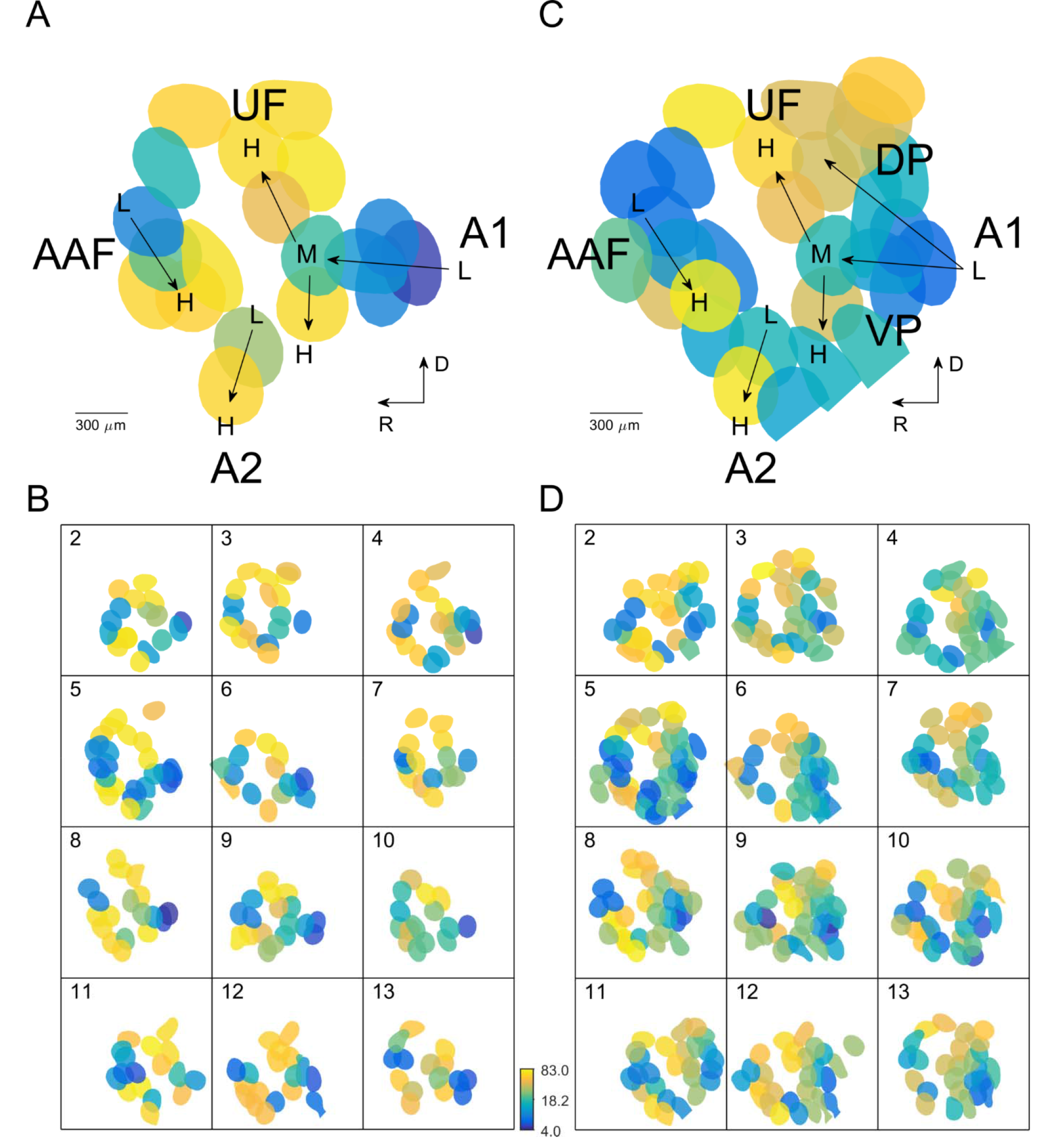
On- and off-response tonotopy. (A) Same example as in Figure 1-4, showing on-response tonotopy. Here, A1, AAF, UF, A2 and DAF ROIs are included as they show prominent on-responses. (B) On-response tonotopy of 12 other mice in this study. (C) Same example as in Figure 1-4, showing off-response tonotopy. In addition to the above-mentioned fields, DP, VP and peri-AAF are also included as they show prominent off-responses. (D) Off-response tonotopy of 12 other mice in this study

We next asked if tonotopy for on- and off-response is robust across single sound levels. We plotted the frequency selectivity at each sound level for both on- and off-response as a function of auditory fields (Figure S4B-F, see Figure S4A for comparison). Indeed, tonotopy was preserved across sound levels for on- and off-responses as manifest by the separation of frequency selectivity into respective low, mid and high frequency band. Together, our results demonstrated that robust tonotopy exists in off-response both at and above threshold.

### Auditory fields have distinct relative on/off-response amplitude with respect to sound level

The tonotopy for on- and off-response indicates a frequency dependency in response amplitude. To investigate the sound-level dependence of on- and off-response amplitude, we summed the response amplitude over frequency for each auditory field (Figure 6). Tones of 20dB SPL did not evoke off-responses while on-response could be seen in certain fields, indicating that off-responses have a higher threshold than on-responses (Figure 6A-D, H-P). However, off-response can have higher amplitude than on-response at highest sound levels (e.g. at 50 and 65dB SPL). We found this to be true in A1 (Figure 6A-C), UF, peri-UF, and posterior-UF (Figure 6D-F). Despite weak on-response, DP and VP also showed larger off-response at high sound levels (Figure 6O, P). The same behavior was not observed in either AAF (Figure 6G-I), A2 (Figure 6L, M), peri-A2 (Figure 6N) or CTR (Figure 6Q). DAF and peri-AAF showed higher off-response amplitude at 65dB SPL (Figure 6J, K), a behavior contrary to the adjacent AAF, validating the rationale to assign these ROIs to separate fields. To quantify the preference of off- versus on-responses across auditory fields we calculated the ratio of off-response versus on-response at 65dB SPL (Figure 6R). DP, UF and peri-UF showed the largest ratios among all auditory fields, thus these areas were preferentially activated by tone offset. Together these results show a differential on/off-response pattern with respect to sound level for different auditory fields. First, in A1, UF and flanking regions such DP and VP, stronger off-response can be observed at 50 and 65dB SPL. Although none of the AAF areas show stronger off-response at higher sound levels, the surrounding peri-AAF and dorsally located DAF show significantly stronger off-response at 65dB SPL. Thus, as one move away from core auditory fields which are strongly activated by tone onset, off-response can be dominant, especially at higher sound levels.

**Figure 6.**
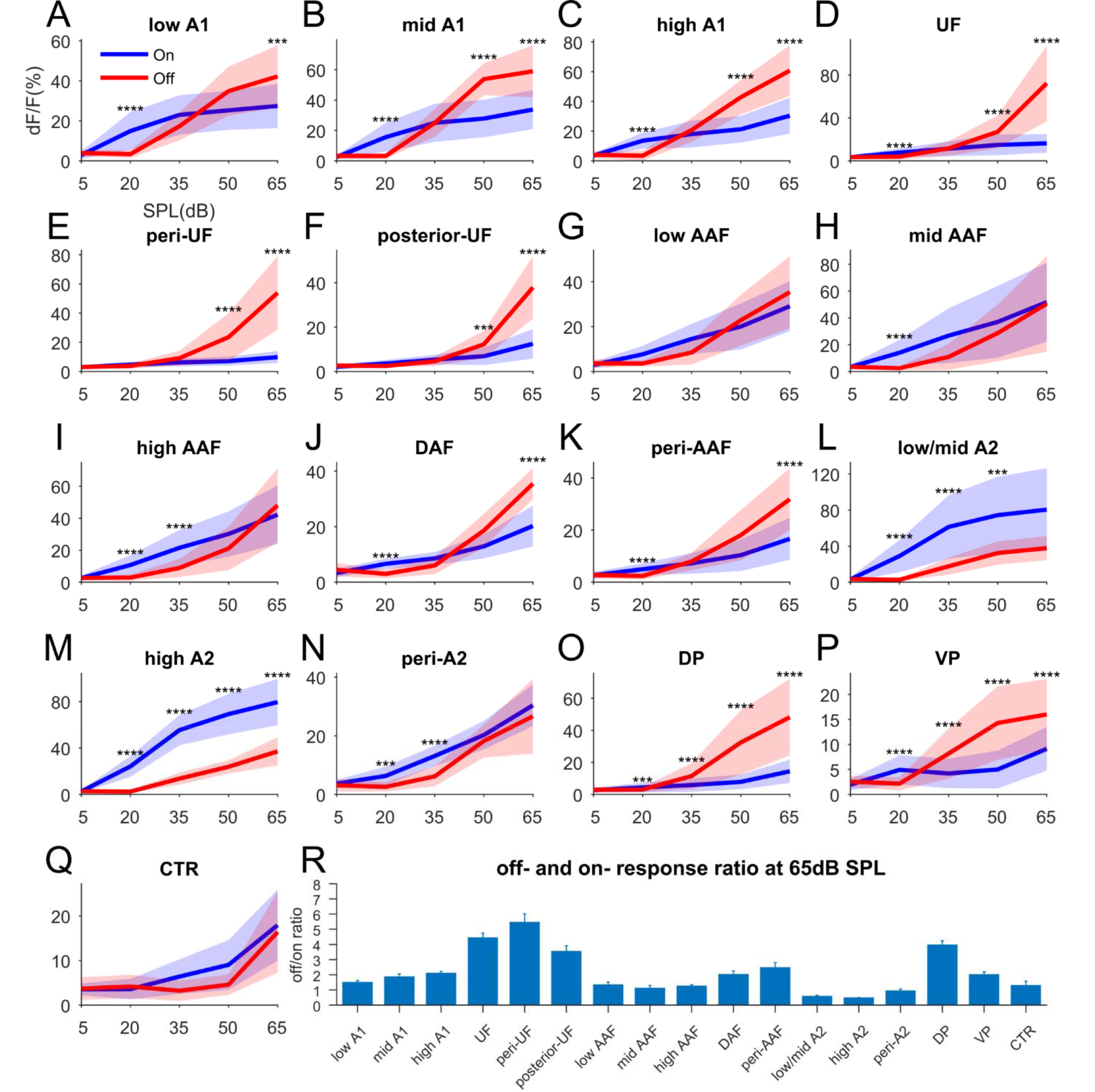
Differential on- and off-response profile as a function of both sound level and auditory fields. (A-Q) On- and off-response profiles with respect to sound level for different auditory fields were obtained by summing over frequency in FRAs. ‘***’ indicates p<0.001; ‘***’ indicates p<0.0001. All shaded regions are SD. (R) Off-and on-response ratio at 65dB SPL. Errorbars show SEM.

### Off-responsive areas are more spatially extensive

Qualitatively the spatial pattern of off-responses appeared more diffusive and elongated in shape than that of on-responses (Figure 1), possibly indicative of underlying circuit differences. To quantify the differences in spatial shape, we first overlaid selected A1, AAF and A2 ROIs with on- and off-response patterns (Figure 7), which serve as landmarks. For on-responses the locations of activation fell into individual ROIs (Figure 7A). However, the areas showing off-responses can span multiple ROIs, which were organized into stripes and were roughly parallel to the dorsal-ventral axis, thus perpendicular to the main rostro-caudal tonotopic axis (Figure 7B). We quantified the extension along the dorsal-ventral axis by computing the signal correlation (SC) among a slice of ROIs that were dorsal to low, mid/high A1 or UF ROIs, respectively. Elongation of off-responsive regions in the dorsal direction should result in a higher off-SCs than on-SCs over distance. Indeed, from low A1, mid/high A1 and UF (Figure 7C), off-SC was significantly higher than on-SC over distance, with mid/high A1 ROIs showing the most prominent extension of high SC values dorsally. Among all the ROIs within field of view, we found that off-SCs were larger than on-SCs at a distance from 0-2mm (Figure 7D), consistent with our finding that areas away from core auditory field such as DP, VP and peri-AAF had prominent off-response. Together these analyses show that off-responses span larger areas than on-responses, and that off-responsive regions in A1 extend dorsally and form elongated spatial patterns perpendicular to the rostrocaudal axis.

**Figure 7.**
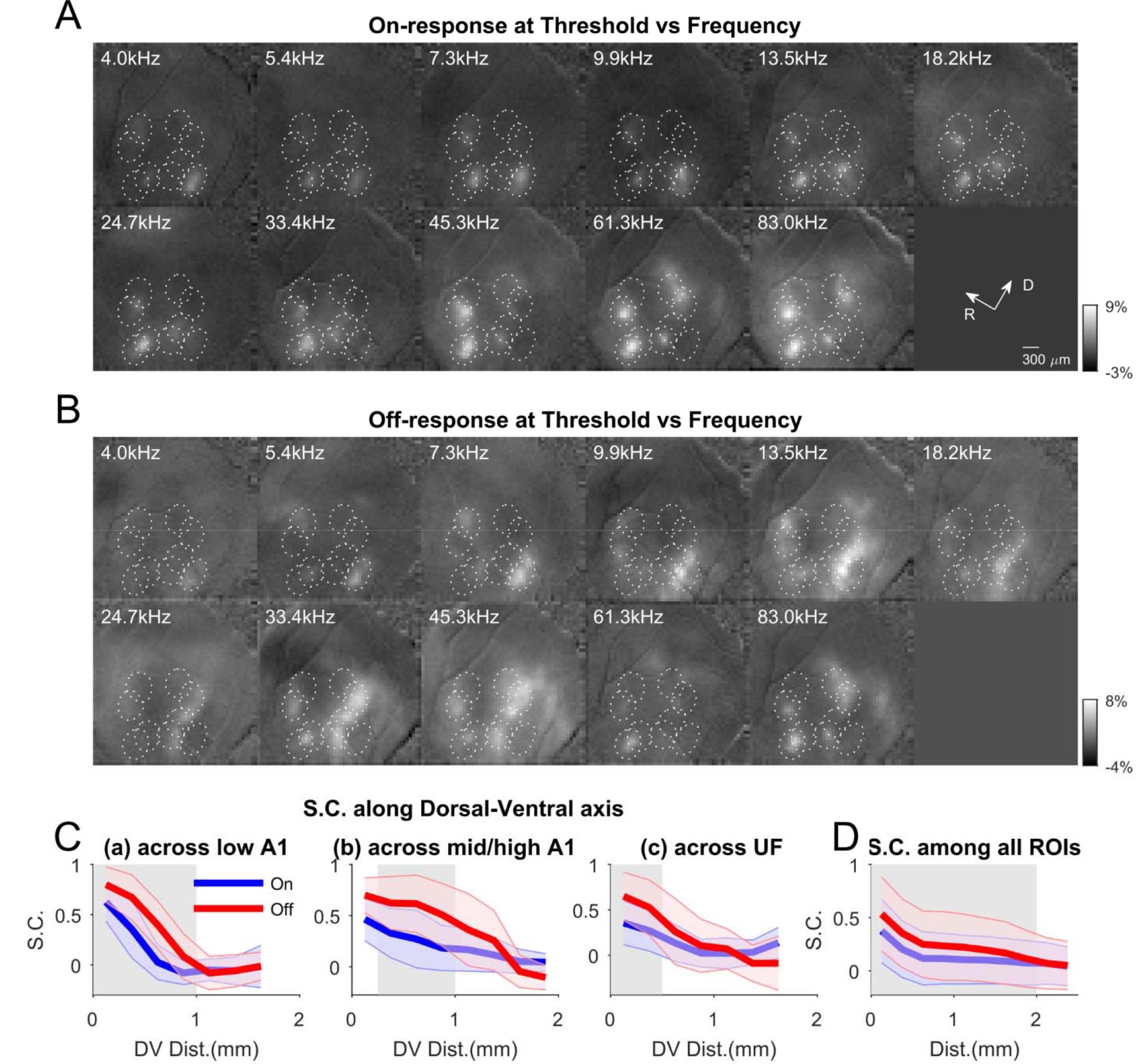
Off-responses are more spatially more extensive and elongated in shape. (A, B) Same example dataset as in Figure 1-4. On- and off-response spatial profiles at respective threshold are plotted as a function of frequency, overlaid with selected A1, UF, AAF and A2 ROIs. Note in (A) local activations tend to fall into individual ROIs while in (B) off-response can span multiple ROIs. (C) On/off signal correlation calculated among ROIs dorsal to low A1 ROIs (a), mid/high A1 ROIs (b), and UF ROIs (c), respectively. (D) On/off signal correlation calculated among all ROis. In (C) and (D) gray region indicates the distance where signal correlation of off-response is higher than that of on-response.

### Different auditory areas exhibit differential combination of dynamics of temporal activation1

So far, we have investigated the spatial pattern of on/off-response but we also noted that different auditory fields can exhibit rich temporal dynamics. To reveal the temporal response patterns, we first determined the distinct temporal activation pattern using k-means clustering of the trial-averaged traces to each stimulus. The dominant subset of seven clusters demonstrated the rich temporal dynamics of the ROIs (Figure 8A, see also Figure S5). The cluster (a)-(e) exhibited different combinations of on/off-response amplitudes. Cluster (e) shows prominent inhibitory effect following the small on-response which might indicate high temporal precision to sound onsets and could enable responses to fast temporally modulated stimuli. In contrast, cluster (f) and (g) showed minimum decay during tone presentation, indicating persistent firing. These results suggest that ACX encode tone duration via three schemes: through firing at onset and/or offset, and through continuous firing.

**Figure 8.**
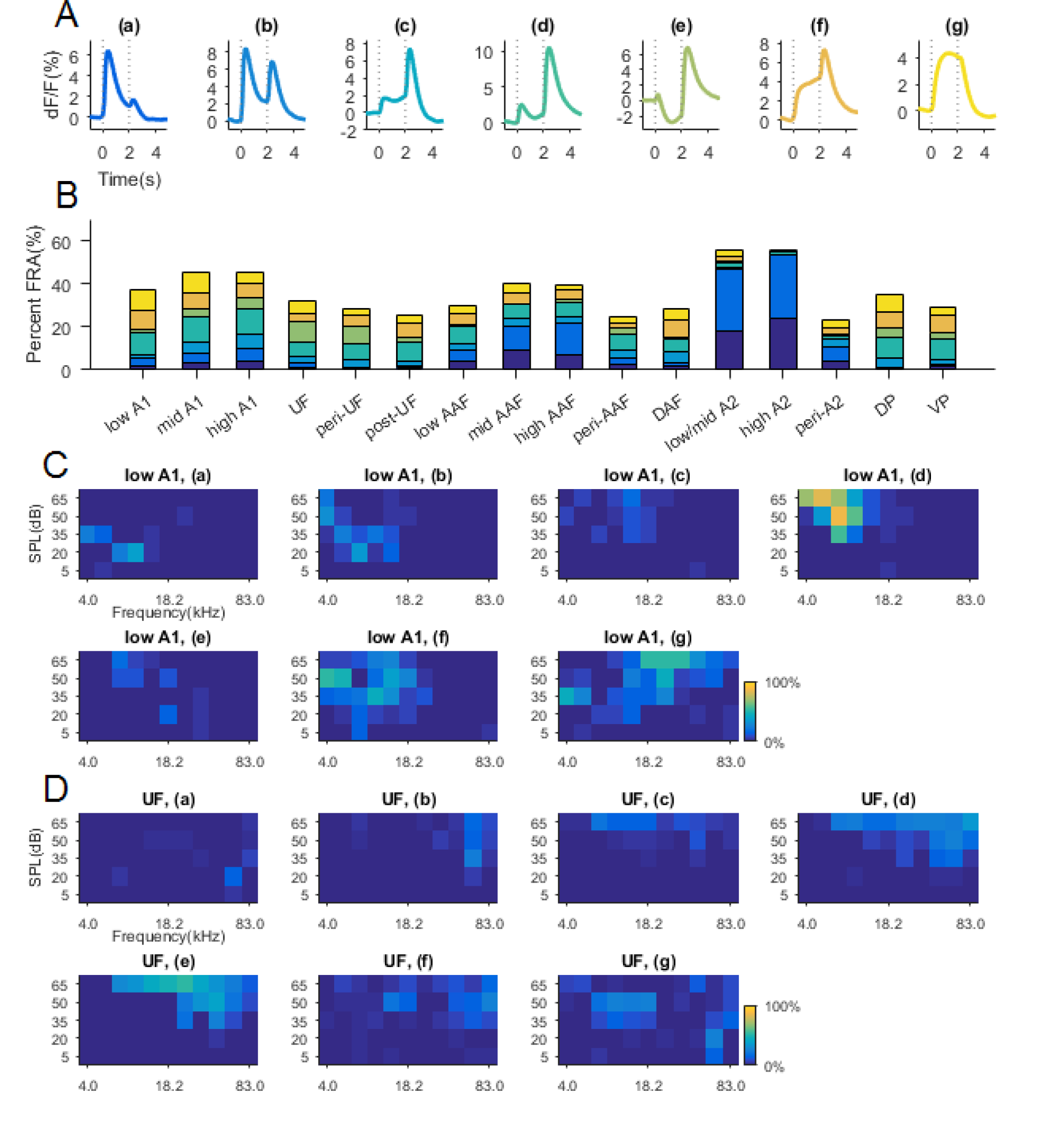
Different temporal dynamics are observed across auditory fields. (A) Mean traces from 7 selected clusters showing typical responses. (B) The proportion of FRA assigned to the 7 clusters in (A) as a function of auditory fields. The same color coding is used in (A) and (B). (C,D) Probability of observing types of temporal traces as a function of frequency and sound level in low A1 (C) and in UF (D). Letter labels correspond to the same cluster labels in (A).

To identify in which ACX fields these temporal dynamics occur, we quantified within each auditory field what percentage of the average traces in response to the different sound stimuli was assigned to each selected cluster (Figure 8B). The fraction of temporal response pattern drawn from each response class varied among fields. For example, while A2 ROIs predominantly show temporal patterns from cluster (a) and (b), A1 ROIs show prominent proportion of traces assigned to cluster (d). In contrast, among all auditory fields, UF and peri-UF had the most proportion assigned to cluster (e), which suggests that these fields might be uniquely sensitive to rapid transients. Low/mid A1, DAF, DP and VP show the most proportion of clusters (f) and (g) among other auditory fields and thus can encode tones with sustained firing. To sum up, different auditory fields have differential combination of temporal dynamics, and some temporal dynamics exist prominently in particular auditory field. This suggests a specialization of the different fields for temporal stimulus attributes.

While largely field-specific, the temporal dynamics could depend on particular stimulus properties. For example, in low A1 most cluster (a) and (b) traces occurred at threshold or close to threshold around low frequency areas (Figure 8C, also see Figure 3B) but cluster (d) traces occurred mostly at high sound level and low frequency combinations. In contrast, cluster (g) traces occurred at complementary (non-BF) regions in FRA, predominantly at edge of FRA, i.e., at mid/high frequency and high sound level combinations. In UF, cluster (a) traces also occurred at high frequency threshold, while cluster (d) and (e) traces occurred predominantly at higher sound level and for mid frequency tones (Figure 8D, see also Figure 3F-H). Cluster (g) traces were found both at high frequency threshold as well as at low/mid frequency and intermediate sound level combinations, and just like low A1, these corresponded to UF’s non-BF regions. Thus, the occurrence of specific type of temporal dynamics is not only a function of auditory field, but can also depend on the particular sound properties.

## Discussion

Temporal information processing is an essential part of auditory system’s function. Here we show that in awake mice, ACX encodes tone offset in a spatially extensive yet tonotopically organized manner with different auditory areas showing distinct selectivity for tone onset or offset. Thus, ACX is spatially organized not only by spectral features, i.e., tonotopy, but also by dynamic features such as sound onset and offset. Moreover, using our machine learning approach we detect areas preferentially activated by tone offset and new auditory areas. Our results show that mouse ACX has a progression of auditory areas from A1 at the caudal pole towards both rostroventral direction (to high frequency A1) and rostrodorsal direction (towards UF), remotely similar to a suggested division of ACX in higher mammals (Rauschecker and Scott, 2009).

While studies in anesthetized mice did not detect tonotopy in off-responses (Baba et al., 2016) we here found robust off-tonotopy across multiple ACX fields, likely due to off-response being most prominent in awake animals (Fishman and Steinschneider, 2009; Joachimsthaler et al., 2014; Qin et al., 2007; Recanzone, 2000). Moreover, we sampled from a broader range of frequencies (4-83.0kHz) and multiple sound levels (5-65dB SPL) which might have aided the detection of tonotopy.

On- and off-responses are thought to be mediated by two non-overlapping set of synapses (Scholl et al., 2010) thus might reflect different inputs to ACX. ACX receives ascending inputs via the lemniscal and non-lemniscal pathways. The lemniscal pathway projects to ACX via the ventral division of the medial geniculate body (MGBv) which shows on-responses (Aitkin and Webster, 1972; Hackett et al., 2011; Imig and Morel, 1983; Redies and Brandner, 1991). Off-responses could potentially parallel this pathway from the superior paraolivary nucleus (SPN) which shows post-inhibition rebound and projects to the inferior colliculus (IC) (Kopp-Scheinpflug et al., 2011), which in turn provides input to MGBv. The non-lemniscal pathway which projects to ACX via the medial and dorsal division of the MGB (MGBm and MGBd) is also a likely source of off-responses. First, off-responses are predominantly observed in a sheet partially surrounding MGBv (He, 2001), while core MGBv mainly shows on-responses. Second, we found that A2 and DP which receive input from MGBd (Lee and Sherman, 2008; Llano and Sherman, 2008) show off-responses. Third, the spatial extensiveness of off-response is consistent with the broad projection from MGBm to ACX (Huang and Winer, 2000; Lee and Winer, 2008). Thus non-lemincal pathways (MGBd and MGBm) likely also provide tone offset information to ACX. Our results show overlapping tonotopy of on- and off-responses albeit areal differences, suggesting that on- and off-response pathways are at least coarsely aligned. Moreover, our results suggest that distinct spatial region in ACX might be formed by differential contributions of the lemniscal and non-lemniscal pathways.

We found that different auditory areas can exhibit distinct temporal dynamics such as a transient increase in firing rate following tone onset and offset, consistent with sparse temporal ACX responses (Hromádka et al., 2008). However, other auditory fields can exhibit other temporal dynamics corresponding to sustained firing throughout tone presentation. Thus, even on a macroscale ACX encode temporal information through either signaling onset and/or offset, or through persistent firing.

To identify ACX regions we employed a novel image segmentation technique to allow us to place ROIs at meaningful locations based on temporal response properties (Whiteway and Butts, 2017). ROIs can be extracted in an unbiased and unsupervised fashion based soly on temporal coativation of pixels, which requires no prior assumptions on the distribution of cortical fields and facilitates identification of known as well as novel areas. Our method captures meaningful variance in image sequence, even though some of the variance was small and yet consistent across trials, which could be easily neglected if analyzed on a pixel by pixel basis. Thus, we provide a general framework which treats cortical fields as individual entities and allows us to study the collective activation as a function of time within each field, providing global information on cortical processing.

Together, our results show that ACX is organized on a macroscale not only based on spectral sound properties but also based on temporal information processing.

## Methods

All procedures were approved by the University of Maryland’s Animal Care and Use Committee.

### Animal

We crossed the CBA/CaJ mice with Thy1-GCaMP6s (JAX stock #024275, GP4.3, (Dana et al., 2014) to obtain F1’s since C57/BL6 are homozygous for Cdh23 allele *ahl*, which causes them to suffer from aging related hearing loss, while CBA/CaJ mice are homozygous for *Ahl+*, which spare them from the phenotype (Kane et al., 2012). F1’s thus have no hearing loss and yet have uniform expression of GCaMP6s under Thy1 promotor in excitatory neurons. We used adult mice of both sexes whose ages range from 2 to 4 months old (female n=7 mice, male n=6 mice).

### Chronic window implant

2-3 hours before surgery, 0.1cc dexamethasone (2mg/ml, VetOne) was injected subcutaneously to reduce brain swelling during craniotomy. Anesthesia was induced with 4% isoflurane (Fluriso, VetOne) with a calibrated vaporizer (Matrx VIP 3000). During surgery, isoflurane level was reduced to and maintained at a level of 1.5%-2%. Body temperature of the animal was maintained at 36.0 degrees Celsius during surgery. Hair on top of head of the animal was removed using Hair Remover Face Cream (Nair), after which Betadine (Purdue Products) and 70% ethanol was applied sequentially 3 times to the surface of the skin before the central part is removed. Soft tissues and muscles were scraped to expose the skull. Then a custom designed 3D printed stainless headplate was mounted over left auditory cortex and secured with C&B-bond (Parkell). A craniotomy with a diameter of about 3.5mm was then performed over left auditory cortex. A three layered cover slip was used as cranial window, which is made by stacking 2 pieces of 3mm coverslips (64-0720 (CS-3R), Warner Instruments) at the center of a 5mm coverslip (64-0700 (CS-5R), Warner Instruments), using optic glue (NOA71, Norland Products). Cranial window was quickly dabbed in kwik-sil (World Precision Instruments) before mounted onto the brain with 3mm coverslips facing down. After kwik-sil cured (2-5min), C&B-bond was applied to secure the cranial window. Synthetic black iron oxide (Alpha Chemicals) was then applied to the hardened surface. 0.05cc Cefazolin (1 gram/vial, West Ward Pharmaceuticals) was injected subcutaneously when entire procedure was finished. After the surgery, the animal was kept warm under heat light for 30 minutes for recovery before returning to home cage. Medicated water (Sulfamethoxazole and Trimethoprim Oral Suspension, USP 200mg/40mg per 5ml, Aurobindo Pharms USA; 6ml solution diluted in 100ml water) substituted normal drinking water for 7 days before any imaging was performed.

### Widefield imaging

Mice were affixed to a custom designed head-post and restrained within a plastic tube. The head of the animal was held upright. Imaging was performed using Ultima-IV two photon microscope (Bruker Technologies) with an orbital nosepiece such that the illuminance light is roughly perpendicular to cranial window (rotation angle was ~60 degrees). As a result, the anterior-posterior axis was not parallel to the edge of the images. 470nm LED light (M470L3, Thorlabs Inc.) was used to excite green fluorescence. Images were acquired with StreamPix 6.5 software (Norpix) at 10Hz and 100ms exposure time. In StreamPix software, we specified the image size to be 400 by 400 with a spatial binning of 3.

### Acoustic stimulus

Pure tones were generated with custom MATLAB script. Each tone lasted 2 seconds with linear ramps of 5ms at the beginning and at the end of the tone. The amplitudes of the tones were calibrated to 75dB with a Brüel & Kjær 4944-A microphone. During sound presentation, sound waveform was loaded into RX6 multi-function processor (Tucker-Davis Technologies (TDT)) and attenuated to desired sound levels by PA5 attenuator (TDT). Then the signal was fed into ED1 speaker driver (TDT), which drove an ES1 electrostatic speaker (TDT). The speaker was placed on the right-hand side of the animal, 10cm away from the head, at an angle of 45 degrees relative to the mid-line. The presentation of tones with various combination of frequencies and sound levels are randomized and controlled by a custom MATLAB program. The silent period in between the 2-second tones was randomly chosen from a uniform distribution between 3 and 3.5 seconds. Frequencies of the tones vary from 4kHz to 83.0kHz with logarithmic spacing and with a density of 16/7 tones per octave. Sound levels vary from 5dB SPL to 65dB SPL with a step of 15dB SPL. Each stimulus was repeated 10 times. In total, the imaging session for each animal lasted ~45min.

### Image preprocessing

We performed three preprocessing steps before using autoencoder for image segmentation. First, we downsampled the original image (400 by 400) using MATLAB (2015b) built-in function wavedec2 (level = 3, wavelet name ‘sym2’, function included in the wavelet toolbox of the same MATLAB version). The resultant image was 52 by 52 in size. The purpose of this step is to both improve signal to noise ratio and to cut the number of image dimensions. Next, we applied a homomorphic filter to correct the non-uniformness in illumination, which was necessary because of the curvature of the brain and given the illumination was often not completely perpendicular to cranial window surface. This preprocessing step removed low-frequency components of the fluorescence as follows. We transformed the downsampled images into Fourier domain, then we multiplied the Fourier transform with a low pass filter such constructed: we first generated a 2D symmetric Gaussian distribution with a standard deviation of 3, and calculated 1 minus the Gaussian distribution. We multiplied the such constructed matrix pointwise with the Fourier transform of the image. The filter would remove the low frequency component in the images and thus correct for illuminance difference across field of view, which is usually slowly varying in space. Finally, we performed an inverse Fourier transform to obtain the desired image. We applied filtering to each image in the image sequence. The third preprocessing step was whitening of the image sequence. We first re-shape each image into column vectors, then we stack them horizontally. Let *I_t_* denote the column vector corresponding to image at time *t*, *M* be the stacked matrix, and *N* be the total number of images:

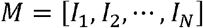

We then subtracted the time average image (<*I*>_t_) from all images:

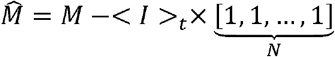

We then performed singular value decomposition on sample covariance matrix of *M̂*:

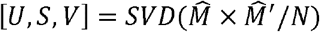

Then we obtained the whitened images using the following equation:

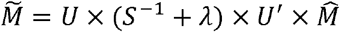

where λ is the regularization term, which we typically picked to be 10^-5^ and we found the behavior of autoencoder to be relatively stable over different choice of λ. We then fed *M̃* into autoencoder algorithm.

### Image Segmentation with constrained autoencoder

We used a dimensionality reduction technique to perform automatic image segmentation such that pixels with strong temporal correlations across the set of images were grouped together into single components (ROIs), following the formulation of Whiteway and Butts (2017). To perform this dimensionality reduction, we used an autoencoder neural network. For each time point *t*, the autoencoder takes the vector of pixel values *y_t_* ∈ ℝ^*N*^ and projects it down onto a lower dimensional space ℝ^*M*^ using an *encoding matrix* W^1^ ∈ ℝ*^M×N^*. A bias term **b**_1_ ∈ ℝ*^M^* is added to this projected vector, so that the resulting vector **z**_t_ ∈ ℝ*^M^* is given by

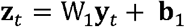

The autoencoder then reconstructs the original activity **y***_t_* by applying a *decoding matrix W_2_* ∈ ℝ*^N×M^* **z***_t_* and adding a bias term **b**_2_ ∈ ℝ^*N*^, so that the reconstructed activity ŷ ∈ ℝ^*N*^ is given by

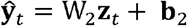

Since the dimensionality of **z***_t_* is typically much smaller than that of **y***_t_*, **z***_t_* should capture variations in **y***_t_* that are shared across many pixels. The entries of W2 then describe how each pixel is related to each dimension of **z***_t_* (see Figure 2C).

The weight matrices and bias terms, grouped as Θ ={W_1_,W_2_,**b**_1_,**b**_2_ }, are simultaneously fit by minimizing the mean square error between the observed activity **y***_t_* and the predicted activity ŷ*_*t*_*:

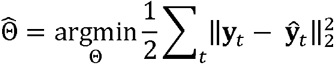

To further enable interpretability of the results, we constrained the weights W_2_ to be non-negative, as one could flip the signs of both spatial and temporal components arbitrarily. This also ensured that all pixels in a given ROI always increase or decrease in intensity together, depending on the sign of **z***_t_*. We also tied the weights such that 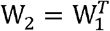. Thus, there was essentially only one spatial weight matrix.

This version of the autoencoder is closely related to principal components analysis (PCA) (Bengio et al., 2013). However, PCA is an inadequate technique for automatic image segmentation since it did not in general result in spatially localized ROIs (Figure S2A), due to the orthogonality constraints imposed by the PCA model. A similar approach to our non-negatively constrained autoencoder is to use non-negative matrix factorization (NNMF) on the preprocessed image sequence. NNMF constrains both the spatial maps and the temporal activations to be non-negative, whereas the RLVM just constrains the spatial maps to be non-negative. The NNMF ROIs also failed to be spatially localized (Figure S2B). Finally, in order to solve the constrained minimization problem above we used the spectral projected gradient method, a constrained variant of gradient descent (Schmidt et al., 2009).

To perform image segmentation with this method we must first specify the number of ROIs (the dimensionality of **z**_*t*_). We determined the appropriate number of ROIs using cross-validation by first fitting the parameters of the autoencoder on 80% of the frames from the image sequence (training data), and then reconstructing the remaining 20% of the images (testing data) using the autoencoder. We then calculated the correlation between the true and reconstructed images on the testing data, as a measurement for goodness of fit. In Figure S1A, we show that with an increasing number of ROIs, the correlation from the testing data increases monotonically, and roughly plateaus after ~50 ROIs. We also performed fitting on the entire image sequence and plot the correlation (Figure S1A, blue curve). A similar monotonic increase is observed, and with 50 or more ROIs, the correlation value is above 0.8, which is quite agreeable considering that the full image sequence consisted of more than 28,000 images. Another criterion we utilized to choose the number of ROIs was the total spatial area covered by the ROIs. An increasing portion of the total area is covered with an increasing number of ROIs, (Figure S1B), and 50 ROIs cover close to 90% of the total area. Given these results, we typically used 50 ROIs in the autoencoder.

To automatically separate isolated areas in single ROIs, e.g., Figure 2C ROI 36, each ROI was first converted to binary image at the threshold of 95 percentile value of the entire image, and then isolated areas were identified using MATLAB built-in function bwlabel. Areas with fewer than 10 pixels were excluded and we would raise this threshold if the final number of total ROIs was more than 1.5 the original number. Isolated areas were also excluded if their summed weights constituted no more than 5% of the total summed weights after thresholding the ROI at the 95 percentile.

### On- and off-response profile

To determine on- and off-response amplitude, first the temporal trace from each trial was normalized to percentage change with respect to baseline fluorescence:

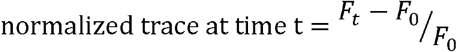

where *F*_0_ is the average of fluorescence within 200ms window before the tone onset. For on-response amplitude, we averaged the normalized trace from 200-500ms after tone onset with the baseline from normalized trace subtracted. For off-response amplitude, we averaged the normalized trace from 200-500ms after tone offset and subtracted the average from the same trace 0-200ms right before tone offset. The 200-500ms window was sufficient to capture the rising phase as well as the peak of the increase in fluorescence in typical on- and off-response. In Figure 3 and Figure S3 all traces plotted were normalized traces obtained in the above-mentioned fashion from each ROI. Widefield FRAs such as shown in Figure 1C, D were also constructed in a same fashion while individual pixels were analyzed.

To obtain on-response frequency profile (Figure 4C-G), we first averaged on-response over repeats and then summed over all sound levels for each ROI. Thus, for each ROI, we obtained one curve for onset frequency response profile. Then we averaged these curves according to the auditory field each ROI was assigned. Similarly, we obtained sound level response profile by summing over frequency (Figure 6). For these two analyses, if the average response over repeats obtained was negative, we would exclude it from the average across ROIs because first, for these analyses we focused on excitatory responses, and second it was indistinguishable if the decrease in fluorescence was a result of inhibition or a natural decay from spontaneous activities.

### Field Parcellation

We assigned ROIs to different auditory fields based upon known tonotopic structure revealed with optical approach (Issa et al., 2014; Tsukano et al., 2015). The general procedure was the same as described in main text. To find peri-AAF, we first identified all the ROIs that were in the vicinity of core AAF. Next, we obtained the summed on-response over all frequencies and sound levels for each ROI, which we refer to as the total on-response. We then sorted the ROIs in descending order based upon the total on-response. Then we summed the total on-response over ROIs and assigned the ROIs whose total on-response summation constitute 80% of the summed total on-response to AAF, while the rest were assigned to peri-AAF. The purpose of doing so was to separate ROIs based on relative strength of total on-response. We defined peri-UF in a similar way. We further separated A1 and AAF ROIs into low, mid and high frequency groups based on the location of the peak in the onset frequency response profile. Frequencies higher than or equal to 4.0kHz but lower than 9.9kHz were considered ‘low’ frequencies. Frequencies higher than or equal to 9.9kHz but lower than or equal to 33.4kHz were considered ‘mid’ frequencies. Frequencies higher than 33.4kHz were considered ‘high’ frequency. We separated A2 ROIs only into low/mid and high frequency group as A2 has a highly-compressed tonotopy gradient.

### On- and off-tonotopy

To establish on- and off-tonotopy, threshold of on- and off-response were first visually determined (Figure 1C, D, white solid lines). For on-tonotopy, tuning curve at threshold was first obtained for each ROI using the on-response identified in the above-mentioned fashion, then we picked 3 frequencies that evoked the largest responses. Among the 3 frequencies, we averaged the 2 frequencies that were closest in number to increase the robustness of our estimate. We then assigned the averaged frequency to the ROI in question as an estimate for the characteristic frequency. We produced the off-tonotopy map in the same fashion but using off-response tuning curve at off-response threshold.

Figure S4 shows the weighted average frequency for different auditory fields. To calculate weighted average frequency, on/off tuning curve was obtained either at threshold or at each sound level, and frequencies were converted to log space before weighted by tuning turve and averaged.

### Signal correlation among ROIs

We used corrected signal correlation (SC) for all our calculation due to the limited number of repeats and the strong tendency of close-by pixels to covary in time (Rothschild et al., 2010; Winkowski and Kanold, 2013). The basic idea is that the uncorrected SC equation contains products of responses from the two ROIs in question on the same trial, and these terms also appear in noise correlation equation. Thus, these products represent to some extent the covariation of ROIs regardless of stimulus presentation, and thus should be excluded from SC calculation. The denominator in the equation was adjusted accordingly to take into account the reduction of number of summation in the nominator.

In Figure 7C(a-c), we calculated SC among selected ROIs that were dorsally located with respect to low A1, mid/high A1 and UF, respectively. To identify these ROIs, we first selected one of the ROIs from, for example, low A1, and then identified all ROIs whose centers were within ~450um to the ROI in question in the rostrocaudal direction but dorsally located. Then we calculated pairwise SCs among all these ROIs and the low A1 ROI, and plotted them as a function of distance.

### Clustering of trace dynamics

Average traces over repeats were obtained from each combination of frequency and sound level. In our study, we used 11 different frequencies and 5 different levels, thus each ROI would contribute 55 different average temporal traces. Each trace had 60 frames with the first 10 frames before sound onset. We pooled these traces across ROIs and across different animals and performed k-means clustering using cosine distance measure because we would like to capture the shape of temporal dynamics regardless of amplitude. To determine the number of proper clusters, we simply used ‘elbow method’ and explored from 5 to 80 clusters, in a step of 5 (Figure S5A). Distortion is the distance of each trace to respective cluster centroid summed over all traces. We picked the number of cluster based on the percentage change of distortion. With 30 clusters, the change in distortion was ~5%, close to our preset threshold. Figure S5B shows all 30 cluster centroids. We picked 7 clusters with the biggest amplitude for further analysis, and they represent typical responses. We quantified the proportion each selected cluster occupies in FRA in different auditory fields by counting the number of average trace assigned to the different types and averaged across ROIs.

## Acknowledgements & Contributions

JL, POK designed research. JL performed experiments and analyzed the data. MW, DB, and JL designed and implemented the analysis algorithm. JL and POK wrote the paper. Supported by NIH RO1DC009607 (POK), NIH T32DC00046 (MRW) and NSF IIS-1350990 (DAB)

